# Dissecting the basis of novel trait evolution in a radiation with widespread phylogenetic discordance

**DOI:** 10.1101/201376

**Authors:** Meng Wu, Jamie L. Kostyun, Matthew W. Hahn, Leonie Moyle

**Affiliations:** Department of Biology, Indiana University, Bloomington, Indiana, U.S.A. 47405; Department of Plant Biology, University of Vermont, Burlington, Vermont, U.S.A., 05401; Department of Computer Science, Indiana University, Bloomington, Indiana, U.S.A. 47405

**Keywords:** phylogenomics, rapid radiation, hemiplasy, *Jaltomata*, *Solanum*

## Abstract

Phylogenetic analyses of trait evolution can provide insight into the evolutionary processes that initiate and drive phenotypic diversification. However, recent phylogenomic studies have revealed extensive gene tree-species tree discordance, which can lead to incorrect inferences of trait evolution if only a single species tree is used for analysis. This phenomenon—dubbed “hemiplasy”—is particularly important to consider during analyses of character evolution in rapidly radiating groups, where discordance is widespread. Here we generate whole-transcriptome data for a phylogenetic analysis of 14 species in the plant genus *Jaltomata* (the sister clade to *Solanum*), which has experienced rapid, recent trait evolution, including in fruit and nectar color, and flower size and shape. Consistent with other radiations, we find evidence for rampant gene tree discordance due to incomplete lineage sorting (ILS) and several introgression events among the well-supported subclades. Since both ILS and introgression increase the probability of hemiplasy, we perform several analyses that take discordance into account while identifying genes that might contribute to phenotypic evolution. Despite discordance, the history of fruit color evolution in *Jaltomata* can be inferred with high confidence, and we find evidence of *de novo* adaptive evolution at individual genes associated with fruit color variation. In contrast, hemiplasy appears to strongly affect inferences about floral character transitions in *Jaltomata*, and we identify candidate loci that could arise either from multiple lineage-specific substitutions or standing ancestral polymorphisms. Our analysis provides a generalizable example of how to manage discordance when identifying loci associated with trait evolution in a radiating lineage.

## INTRODUCTION

Phylogenies contribute to our understanding of the evolutionary history of traits (Felsenstein, 1985). When the patterns of relationship among species is known, robust inferences about character state evolution can be made, including the number of times a character evolved, the direction of character evolution, and the most likely ancestral character state. Phylogenies can also reveal whether lineages with similar phenotypic traits have evolved these via independent evolution (convergence or parallelism) or whether a single origin is more likely (Wake et al., 2011). The recent use of whole genomes or transcriptomes to make phylogenetic inferences from thousands to millions of sites (“phylogenomics”), has succeeded in its aim of generating species trees with high levels of statistical support. However, other genome-wide analyses have begun to reveal unexpected complexities in the evolutionary history of rapidly radiating lineages—including widespread gene tree discordance due to incomplete lineage sorting and/or introgression (Degnan & Rosenberg, 2009). This frequent discordance among individual gene trees can amplify incorrect inferences of trait evolution on even well-supported species trees. In particular, when a trait is determined by genes whose topologies do not match the species topology, incorrect inferences of homoplasy (independent evolution of the same character state) are substantially elevated—a phenomenon known as ‘hemiplasy’ (Avise & Robinson, 2008; Hahn & Nakhleh, 2016; Storz, 2016). Because understanding trait evolution—including the underlying genetic changes—is of particular interest in species radiations, extra care must be taken to consider and account for the influence of hemiplasy in these cases.

The fraction of the genome affected by hemiplasy will depend upon the amount and sources of gene tree discordance in a clade. In rapidly radiating species groups, widespread discordance has been attributed to the effects of both incomplete lineage sorting (ILS) and introgression between lineages (Degnan & Rosenberg, 2009). ILS affects gene tree topologies when segregating ancestral variation is maintained through consecutive speciation events (Maddison, 1997). Because the effect of ILS is proportional to ancestral population size, and inversely proportional to the time between speciation events (Pamilo & Nei, 1988), ILS is expected to be particularly exaggerated in radiations where a diverse ancestral population undergoes rapid speciation. Indeed, gene tree discordance has been noted for a substantial fraction of the genome in rapidly radiating groups, including the *Drosophila simulans* sub-clade (Garrigan et al., 2012), African cichlid fishes (Brawand et al., 2014), wild tomatoes (Pease et al., 2016), and the genus *Arabidopsis* (Novikova et al., 2016). When there is introgression, discordance emerges because genes that are introgressed among lineages will show historical patterns of relatedness that differ from the loci in the genome into which they are introduced. Substantial introgression has also been identified among rapidly radiating lineages through genome-wide analysis, including in *Xiphophorus* fishes (Cui et al., 2013), *Heliconius* butterflies (Martin et al., 2013), Darwin’s finches (Lamichhaney et al., 2015), and *Anopheles* mosquitoes (Fontaine et al., 2015).

Both ILS and introgression contribute to hemiplasy because they cause a proportion of gene trees to disagree with the species tree (Avise & Robinson, 2008; Hahn & Nakhleh, 2016; Storz, 2016). Specifically, the probability of hemiplasy is expected to be 1) proportional to the fraction of gene trees that are discordant with the species tree; and 2) negatively correlated with the branch length leading to clades with similar phenotypes (Hahn & Nakhleh, 2016). A higher proportion of discordant gene trees increases the probability that a character of interest is underpinned by genes that have a tree topology that differs from the species tree; shorter branch lengths increase the chance of incorrectly inferring homoplasy, as they leave relatively little time for convergent evolution to happen (Hahn & Nakhleh, 2016). Both conditions are expected to be exaggerated specifically in rapidly diversifying groups. Therefore, in these cases mapping characters onto a single species tree has a substantially elevated risk of incorrectly inferring the number of times a trait has evolved and the timing of trait changes (Avise & Robinson, 2008; Hahn & Nakhleh, 2016; Storz, 2016). Hemiplasy also affects inferences about the specific loci inferred to underlie trait transitions because, when ILS or introgression are common, the substitutions underlying trait transitions may occur on gene trees that are discordant with the species tree (Mendes et al., 2016). Accordingly, genome-wide analyses must take into account the extent and distribution of ILS and introgression if they are to accurately infer the number and timing of evolutionary changes in specific traits, and the genes underlying these changes.

In this study, we used genome-wide data to investigate the morphologically and ecologically diverse plant genus *Jaltomata*, in which several key trait transitions appear to have occurred in parallel (Miller et al., 2011), and have been inferred to be independent convergent responses to similar selective pressures. However, because trait diversification has occurred in a relatively short period in this group, the probability of hemiplasy is also expected to be elevated. Our main goals were to assess the timing of lineage and trait diversification in the group, and to identify sources of genetic variation that potentially contribute to rapid trait diversification in *Jaltomata*, while taking into account the potential for hemiplasy. To do so, we generated a clade-wide whole-transcriptome dataset and explicitly evaluated alternative scenarios to explain trait evolution by: 1) reconstructing phylogenetic relationships among target species, and evaluating the extent of discordance with the resulting inferred species tree; 2) evaluating patterns of trait variation and evolution in key reproductive (flower and fruit) characters, in the context of best and least supported nodes in this tree; and, 3) evaluating specific scenarios of the genetic changes associated with this trait evolution, in order to identify candidate loci that might be causally responsible. Our results imply two different scenarios of trait evolution for fruit color versus floral traits, reflecting the different amounts of hemiplasy associated with the two traits. While fruit color evolution in *Jaltomata* could be confidently inferred—along with potential *de novo* molecular changes on the relevant evolutionary branches—inferring the history of floral trait evolution and the potential contributing loci requires more careful treatment that considers the high probability of hemiplasy.

## MATERIALS AND METHODS

### Study system

The plant genus *Jaltomata* includes approximately 60-80 species, distributed from the southwestern United States through to the Andes of South America (Mione, 1992; Mione et al., 2015) (Figure 1). It is the sister genus to *Solanum*, the largest and most economically important genus in the family Solanaceae (Olmstead et al., 2008; Särkinen et al., 2013). Species of *Jaltomata* live in a wide range of habitats, and are phenotypically diverse in vegetative, floral, and other reproductive traits (Mione 1992; Kostyun & Moyle 2017). Floral diversity is particularly pronounced in *Jaltomata*. In comparison to closely related clades (including *Solanum, Capsicum*, and *Lycianthes*) which predominately have ‘flattened’ rotate corollas (petals) (Knapp, 2010), Jaltomata species exhibit a variety of corolla shapes, including rotate, as well as campanulate and tubular (Miller et al., 2011). All *Jaltomata* species also produce at least some nectar, including noticeably red- or orange-colored nectar in some lineages, while nectar is not produced by species in *Solanum*.

**Fig 1.**
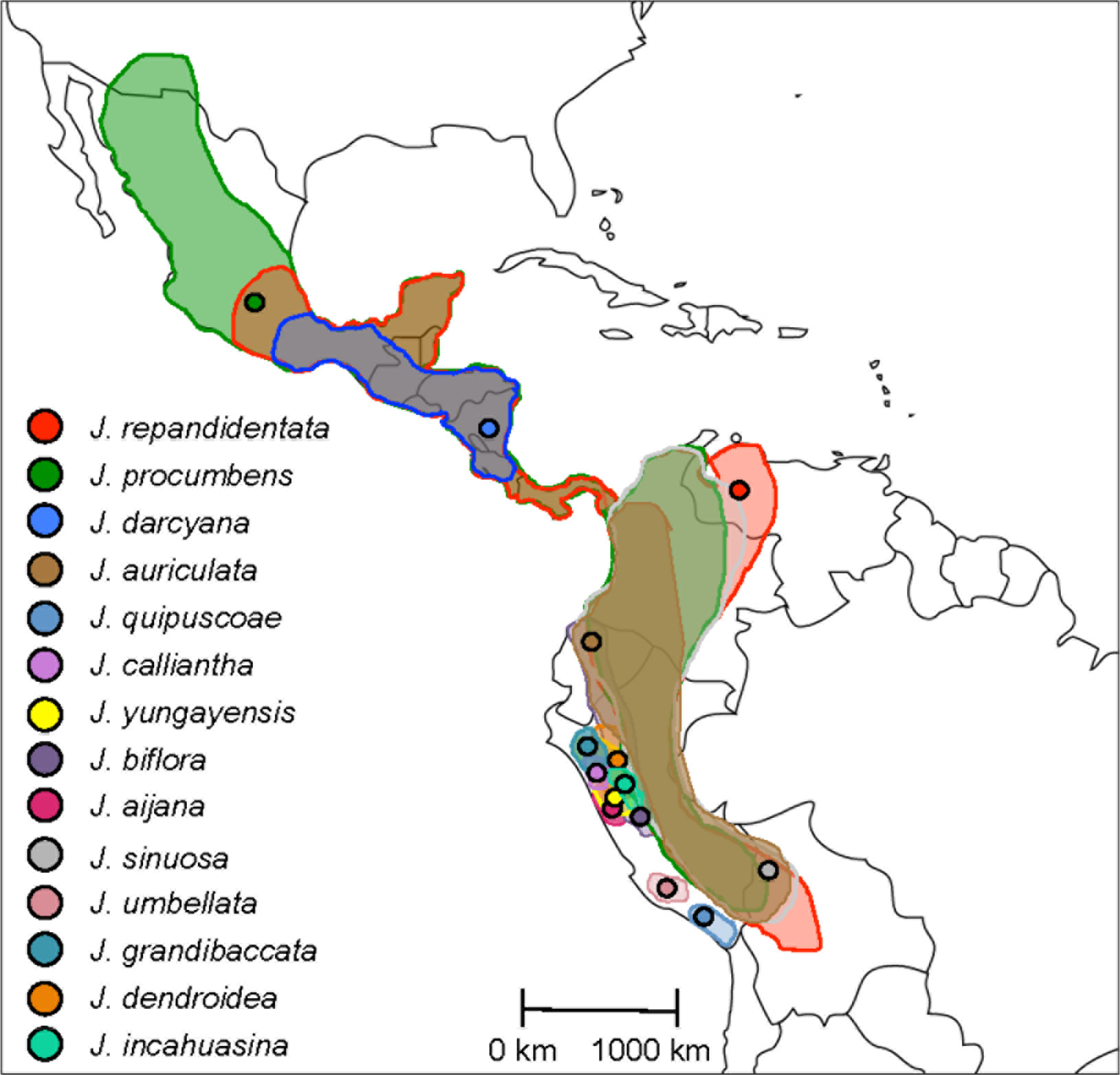
Geographic distribution of investigated *Jaltomata* species. For each species, the sample location is labeled. Ranges estimated from herbarium specimens (T. Mione and S. Leiva G., pers. comm.; J. L. Kostyun, unpub.).

Species also differ in fruit color, and fruit color variation appears to characterize major subgroups within the genus as separate dark purple, red, and orange-fruited clades (Miller et al., 2011; Särkinen et al., 2013). Several species also have green fruit at maturity, although these lineages appear to be distributed across the three major *Jaltomata* clades, suggesting multiple convergent losses of fruit pigment (Miller et al., 2011). The first molecular phylogeny of this genus (Miller et al., 2011) was inferred from a single gene (*waxy*), and indicates that the lineage of species with red fruits is sister to the rest of the genus. However, a more recent study using seven loci (5 plastid and 2 nuclear, including *waxy*) showed a conflicting topology, with purple-fruited lineages sister to the remaining groups, and red-fruited lineages more closely related to lineages with orange fruits (Särkinen et al., 2013). The inconsistency between the two studies might be the result of using few loci, or of reconstructions performed with loci that have different evolutionary histories.

### RNA preparation, sequencing, and transcript assembly

We chose 14 target *Jaltomata* species that are distributed across the three previously identified major clades (Miller et al., 2011), and that span representative floral diversity within the genus (Figure 2A, Table S1). Tissues for RNA extraction included seven reproductive tissues (ranging from early bud, to mature pollinated flower, to early fruit) and four vegetative tissues (roots, early leaf buds, and young and mature leaves), from a single representative individual of each target species (see Supplementary text). All sampled individuals were housed at the Indiana University research greenhouse, under standardized temperature (15-20°C), watering (twice daily), and lighting (16-hour days) conditions.

**Fig 2.**
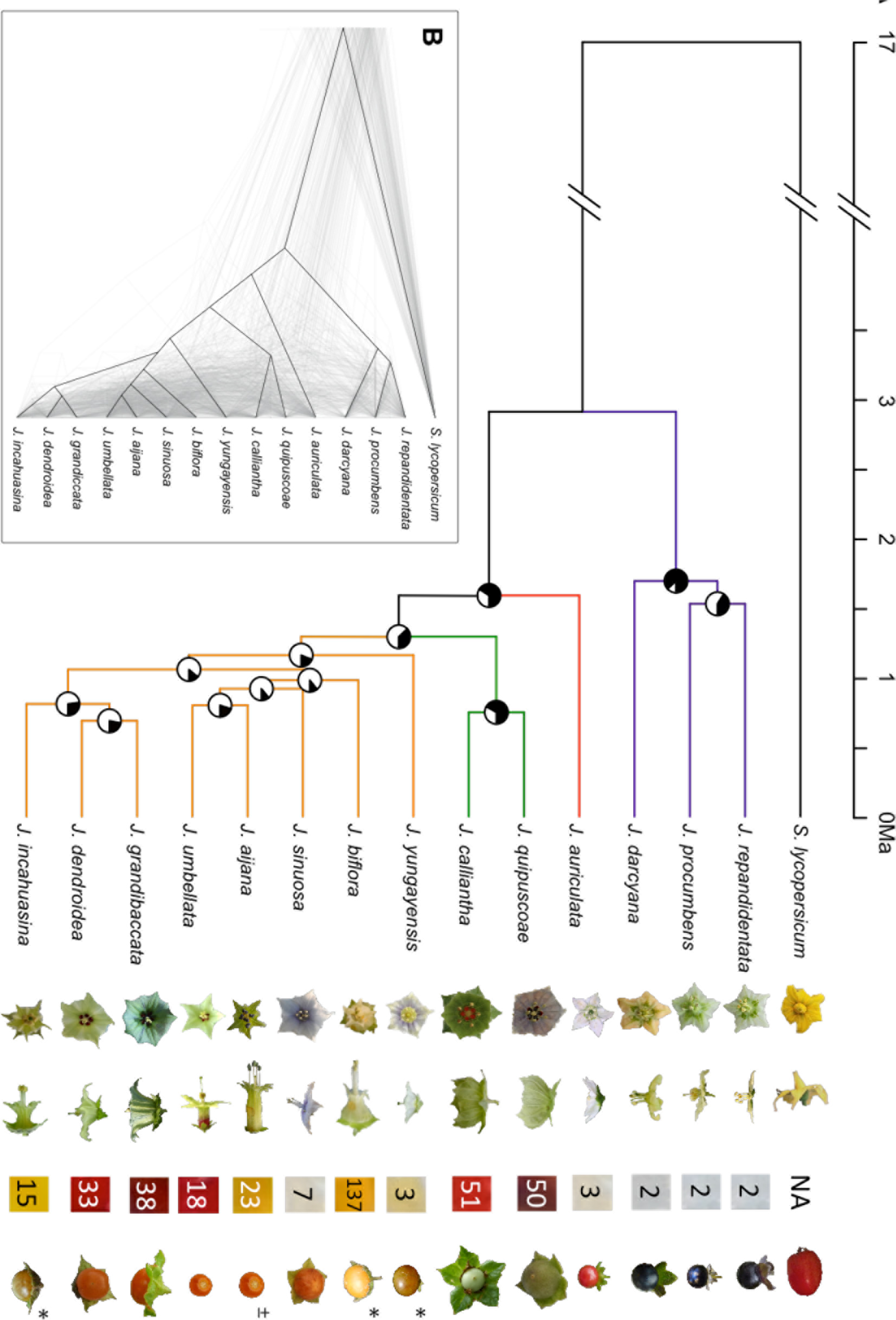
The phylogeny of investigated *Jaltomata* species. **(A)** A whole-transcriptome concatenated phylogeny of *Jaltomata* species with *Solanum lycopersicum* as outgroup. The Pie chart on each internode shows the concordance factor estimated from BUCKy, with the amount of black representing the degree of concordance. Divergence times estimated in *ape* with 17MYA *Jaltomata-Solanum* calibration from (Särkinen et al., 2013). Representative flower and fruit images to the right of species names: front view of flower, lateral view of flower, nectar color and volume (uL) per flower, and ripe fruit. Image with ± indicates that fruit from a similar species is shown, and * indicates contributed by Dr. Thomas Mione at Central Connecticut State University. **(B)** A ‘cloudogram’ of 183 gene trees whose average bootstrap values are larger than 70 across the nodes. For contrast, the concatenated tree (Fig 2A) is shown in black.

Tissue collection and RNA extraction followed Pease et al. (2016): briefly, tissue was collected into pre-chilled tubes under liquid nitrogen, each sample was individually ground under liquid nitrogen, and RNA was extracted from <100mg ground tissue using the Qiagen Plant RNeasy kit. RNA quality/quantity was checked via Nanodrop (Thermo Fisher Scientific); qualified samples of >50ng/uL with 260/280 and 260/230 between 1.8-2.0 were brought to the IU Center for Genomics and Bioinformatics (CGB) for library preparation. Separate reproductive and vegetative libraries for RNA-seq were prepared by pooling equi-molar RNA samples from all reproductive tissues, and all vegetative tissues, respectively, for each species. Both reproductive and vegetative libraries were prepared for all species except for *J*. *grandibaccata*, for which only vegetative RNA could be obtained.

Libraries were sequenced using 100-bp paired-end reads in a single lane of Illumina Hi-seq 2000 (San Diego, CA, USA). Raw paired-end reads were filtered for quality using the program Shear (https://github.com/jbpease/shear) by removing low quality reads and ambiguous bases, and trimming adapter ends (see Supplementary text). The retained reads (length >50 bps) from vegetative and reproductive transcriptomes of the same species were combined prior to assembly using Trinity with the default settings (Grabherr et al., 2011). The open reading frame of each assembled transcript was predicted using TransDecoder v.2.0.1 with default settings (Haas et al., 2013). All the predicted protein-coding sequences within each *Jaltomata* species transcriptome were reduced using CD-HIT v4.6 with -c 0.99 -n 10 (Fu et al., 2012). Each sequence in the assembled transcriptome was presented as a haploid representative of particular transcript. To include domestic tomato (*Solanum lycopersicum*) as the outgroup in the following analyses, we also downloaded the annotated tomato protein-coding sequences from SolGenomics (ftp://ftp.solgenomic.net).

### Protein-coding gene ortholog identification

To infer orthologous gene clusters, we followed a pipeline designed for transcriptome data in non-model species, that begins with an all-by-all BLAST search followed by several steps that iteratively split sub-clusters of homologs at long internal branches, until the subtree with the highest number of non-repeating/non-redundant taxa is obtained (Yang et al., 2015; Yang & Smith, 2014) (see Supplementary text, Figure S1). For the primary analyses, our homologous clusters were required to include a *S*. *lycopersicum* (tomato) homolog in each cluster. For one of our downstream analyses (molecular evolution on the basal branch leading to *Jaltomata*; see below) we also used *Capsicum annuum* (pepper) sequence data. To do so, we added the *C*. *annuum* sequences if the tomato sequence in the orthologous cluster had an identified 1-to-1 ortholog in a pepper gene model (http://peppersequence.genomics.cn).

We prepared multiple sequence alignments of orthologous genes using the program GUIDANCE v.2.0 (Sela et al., 2015) with PRANK v.150803 (Löytynoja & Goldman, 2005) as the alignment algorithm, with codons enforced and ten bootstrap replicates. As a final quality check, we further removed poorly aligned regions using a sliding window approach that masked any 15-bp window from alignment if it had more than three mismatches (indels/gaps were not counted) between ingroup sequences, or had more than five/seven mismatches when tomato/pepper sequences were included. After this process, any alignment with more than 20% of its sequence masked was removed from the analysis. The resulting sequence alignments were converted to the Multisample Variant Format (MVF), and then genetic distances were computed in all possible pairs of species using the program MVFtools (Pease & Rosenzweig, 2015).

### Estimating the amount of shared variation

To quantify the amount of variation shared among species and subclades in *Jaltomata*, the reads from all 14 species were mapped to the reference tomato genome (The Tomato Genome Consortium, 2012) using STAR v2.5.2 (Dobin et al., 2013). SAM files generated were converted to sorted BAM files using SAMtools v. 0.1.19 (Li et al., 2009). SAMtools *mpileup* was then used to call alleles from the BAM files for all lineages. VCF files were processed into MVF files using *vcf2mvf* from the MVFtools package (Pease & Rosenzweig, 2015), requiring non-reference allele calls to have Phred scores ≥ 30 and mapped read coverage ≥ 10. Based on the MVF files, the numbers of variant sites shared between different subclades of *Jaltomata* species were counted.

### Phylogenetic analysis

We used four different, but complementary, inference approaches to perform phylogenetic reconstruction: 1) maximum likelihood applied to concatenated alignments; 2) consensus of gene trees; 3) quartet-based gene tree reconciliation; and 4) Bayesian concordance of gene trees. Because these four approaches use different methods to generate a phylogeny, we applied all four to evaluate the extent to which they generated phylogenies that disagreed, as well as to identify the specific nodes and branches that were robust to all methods of phylogenetic reconstruction. For the concatenation approach, we first aligned all orthologous genes (n=6431), and then used those alignments to build a supermatrix of sequences (6,223,350 sites in total). The species tree was then inferred by maximum likelihood using the GTRCAT model in RAxML v8.23 with 100 bootstraps (Stamatakis, 2006). We also inferred chromosome-concatenated phylogenies with this method. The other three methods (i.e. consensus, quartet-based, and Bayesian concordance) infer species relationships based on gene trees. First, we inferred the majority rule consensus tree with internode certainty (IC) and internode certainty all (ICA) support scores using RAxML with the option for Majority Rule Extended (Salichos & Rokas, 2013). Second, we inferred a quartet-based estimation of the species tree by using the program ASTRAL v.4.10.9 with 100 bootstraps (Mirarab & Warnow, 2015). All 6431 RAxML gene trees were used as input in the consensus and quartet-based approaches. Finally, the Bayesian primary concordance tree and associated concordance factors (CFs; indicative of the posterior probability of gene trees supporting a node) at each internode of the primary concordance tree was computed in the program BUCKy v1.4.4 (Larget et al., 2010). Because BUCKy is computationally intensive for a large number of input gene trees, we only used orthologous gene sets that were potentially informative for resolving gene trees in these analyses. Specifically, we utilized 1517 genes that showed average bootstrap values >50 across the RAxML-inferred gene trees. (These are generally the loci with sufficient genetic variation across the tree to provide information about branch support.) The input of a posterior distribution of gene trees was generated from an analysis with MrBayes v3.2 (Huelsenbeck & Ronquist, 2001). We ran MrBayes for one million Markov chain Monte Carlo (MCMC) generations, and every 1000th tree was sampled. After discarding the first half of the 1000 resulting trees from MrBayes as burnin, BUCKy was performed for one million generations with the default prior probability that two randomly sampled gene trees share the same tree topology is 50% (α = 1) (Larget et al., 2010).

All inferred species trees were plotted using the R package “phytools” (Revell, 2012). To estimate dates of divergence, we used the function “chronos” in the R package “ape” (Paradis et al., 2004) to fit a chronogram to the RAxML genome-wide concatenated phylogeny by using penalized likelihood and maximum likelihood methods implemented in chronos. Times were calibrated using a previous estimate of the divergence time between *Solanum* and *Jaltomata* at ~17 Ma (Särkinen et al., 2013). To visualize gene tree discordance, a “cloudogram” of 183 gene trees with average node bootstrap values greater than 70 was prepared using DensiTree v 2.2.1 (Bouckaert, 2010).

### Ancestral state reconstruction

The number of sampled species (14) is small compared to the size of this clade (60-80 species), and sparse taxon sampling is known to affect the reconstruction of ancestral character states (Heath et al., 2008). Nonetheless, to assess whether our confidence in the number and placement of transitions generally differs among different traits in our clade, we reconstructed ancestral states for fruit color, nectar color, nectar volume and corolla shape. We used the ultrametric species tree inferred from the RAxML genome-wide concatenated phylogeny and the distribution of traits at the tips of phylogeny (Figure 2A) as input. While nectar volume is a quantitative trait, the other three traits are categorical (fruit color: purple/red/orange/green; nectar color: red/clear; corolla shape: rotate/campanulate/tubular). Ancestral character states were inferred using the standard maximum likelihood method with the equal rates model “ER” within the phytools package (Revell, 2012), which models the evolution of discrete-valued traits using a Markov chain, and the evolution of continuous-valued traits using Brownian motion.

### Testing for introgression

We searched for evidence of post-speciation gene flow, or introgression, using the ABBA-BABA test (Durand et al., 2011; Green et al., 2010) on the concatenated orthologous sequence alignment (n=6431 and 6,223,350 sites in total). The ABBA-BABA test detects introgression by comparing the frequency of alternate ancestral (“A”) and derived (“B”) allele patterns among four taxa. In the absence of gene flow, the alternate patterns ABBA and BABA should be approximately equally frequent, given the equal chance of either coalescence pattern under incomplete lineage sorting (ILS). An excess of either ABBA or BABA patterns is indicative of gene flow. Because of the low resolution of many recent branches within major clades of the phylogenetic tree (see Results), evidence for introgression was only evaluated between four well-supported major subclades, each of which is characterized by a distinct fruit color in our specific dataset (i.e. the purple-, red-, and orange-fruited major clades, and a two-species clade of green-fruited taxa; see Results). Patterson’s *D*-statistic was calculated for all four-taxon combinations including one taxon from the green-fruit lineage, one from red or orange-fruit lineage, one from purple-fruit lineages, and one from tomato as the outgroup. Patterson’s *D*-statistic is calculated as (ABBA-BABA) / (ABBA+BABA) for biallelic sites in the multiple sequence alignment (Durand et al., 2011; Green et al., 2010). To further investigate the taxa involved in and the direction of introgression specifically involving purple-fruited lineages, we also used a symmetric five-taxon phylogeny method ‘*D*-foil’ test (Pease & Hahn, 2015) on the transcriptome-wide concatenated dataset, however the results of these analyses were inconclusive (see Supplementary text).

### Identifying genetic variation associated with trait evolution

We used two general strategies to identify loci that might contribute to important phenotypic trait (fruit and floral) transitions within *Jaltomata*. First, to identify loci that have experienced lineage-specific *de novo* adaptive molecular evolution, we evaluated loci for patterns of molecular evolution indicative of positive selection on specific phylogenetic branches (i.e. *d*_*N*_/*d*_*S*_ > 1). Second, to identify variants that might have been selected from segregating ancestral variation, we identified genetic variants that had polyphyletic topologies that grouped lineages according to shared trait variation rather than phylogenetic relationships (‘PhyloGWAS’ Pease et al., 2016).

*1) Lineage-specific* de novo *evolution associated with trait variation:* We identified loci with signatures of *de novo* adaptive molecular evolution (i.e. significantly elevated rates of non-synonymous substitution) across each available locus in our transcriptome (sometimes called ‘reverse ecology’ Li et al., 2008) as well as in a set of *a priori* candidate loci identified based on known or putative functional roles associated with floral or fruit trait variation (Krizek & Anderson, 2013; Rausher, 2008; Specht & Howarth, 2015) (see supplementary text). Tests were only performed on the four best-supported branches within the phylogeny (see Results). For each locus (group of orthologs), we inferred putative adaptive evolution (i.e. *d*_*N*_/*d*_*S*_ > 1) using PAML v4.4 branch-site model (model = 2 and NS sites = 2) on the target branches (Yang, 2007). In each analysis, a likelihood ratio test (LRT) was used to determine whether the alternative test model (fixed_omega = 0) was significantly better than the null model (fixed_omega = 1). In addition, because PAML uses a tree-based *d*_*N*_/*d*_*S*_ model to reconstruct ancestral states and lineage-specific substitutions,_and because high levels of incongruence of gene trees caused by ILS and introgression can produce misleading results when gene trees do not match the assumed species tree (Mendes et al., 2016; Pease et al., 2016), we limited our tests of molecular evolution to the subset of genes for which 1) the RAxML gene tree contained the target ancestral branch (that is, the target branch was supported by the genealogy of the tested/target locus); and 2) there was at least one non-synonymous substitution that could be unambiguously assigned to this branch.

Prior to testing individual loci, we further filtered our data to ensure that poorly aligned and/or error-rich regions were excluded from our alignments (as these tests are particularly sensitive to alignment errors that generate spurious non-synonymous changes). To do so, we used the program SWAMP v1.0 to remove regions from alignments when they showed higher than expected non-synonymous substitutions (i.e. more than five non-synonymous substitutions in 15 codons; the second sliding-window sequence alignment check, different from above) and a minimum sequence length of 50 codons (Harrison et al., 2014). We also required that each alignment must contain a tomato sequence and orthologous sequences from all investigated *Jaltomata* species (except for *J*. *grandibaccata* because reproductive tissues were not sampled from this species)._The resulting sequence alignments were converted to codon-based MVF file format (Pease & Rosenzweig, 2015), prior to performing branch-site tests. We first defined putative genes showing positive selection by using the uncorrected *p*-value <0.01 as cutoff. The false discovery rate (FDR; (Benjamini & Hochberg, 1995) was then calculated for the PAML *p*-values in each branch-specific test.

For the PAML analyses of *a priori* candidates, we used a slightly less stringent statistical cutoff for consideration and required only that each cluster of orthologous sequences had a minimum representation of species from each of the major clades (see supplementary text); this allowed us to evaluate more of these loci while still testing molecular evolution only on the four well-supported branches. For *a priori* genes showing a significant signature of positive selection (*p* <0.05) and for genes identified by the genome-wide unbiased analyses (FDR <0.1), we manually checked the sequence alignments to examine whether they contain putative multi-nucleotide mutations (MNMs), which can cause false inferences of positive selection in the PAML branch-site test (Venkat et al., 2017). Here we assigned an MNM in cases where we observed that a single codon had 2 or 3 substitutions on the selected branch.

To determine the putative functional categories of genes with elevated per site non-synonymous substitution rates, and to assess whether these were enriched for particular functional categories, genes with uncorrected *p*-value <0.05 from the transcriptome-wide analysis were also examined using Gene Ontology (GO) terms. GO term reference was obtained from the Gene Ontology project (www.geneontology.org). GO terms for each gene were obtained from SolGenomics (ftp://www.solgenomics.net). GO term enrichment analysis was performed with ONTOLOGIZER v2.0 using the parent-child analysis (Bauer et al., 2008).

*2) Ancestral genetic variation associated with trait variation:* To identify shared ancestral variants that were associated with trait variation across lineages, we used a ‘PhyloGWAS’ approach (Pease et al., 2016) in which we searched for SNPs that were shared by current accessions that share the same character state, regardless of their phylogenetic relatedness. This approach is only informative in cases where trait variation is not confounded with phylogenetic relationships, which in *Jaltomata* applies to floral shape transitions from the ancestral ‘rotate’ form to the two derived forms (see Results). For our analysis, we treated both campanulate and tubular corolla as the derived state, and rotate corolla as the ancestral state (Figure S2). These categories of floral shape are also perfectly associated with nectar color variation; species with rotate corollas have small amounts of clear/very lightly colored nectar (ancestral), whereas species with campanulate or tubular corollas have larger amounts of darkly colored red or orange nectar (derived). To assess whether the number of nonsynonymous variants found to be associated with our defined groups of floral traits (see Results) was greater than expected by chance, we generated a null distribution due to ILS alone by simulating datasets over the species tree (Figure 2A) using the program *ms* (Hudson, 2002). An associated *p* value was determined by the proportion of simulated datasets that have a greater number of genes perfectly associated with the floral trait distribution than our observed value (see Supplementary text).

## RESULTS

### Transcriptome assembly and ortholog inference identified >6000 orthologs

In assembled transcriptomes from both reproductive and vegetative tissues for each of 14 *Jaltomata* species (except for *J*. *grandibaccata*, which only included vegetative tissues), the number of transcripts per lineage ranged from 46,841-132,050, and mean transcript length ranged from 736-925 bp (Table S2). Based on our criteria for ortholog identification (see Methods, Figure S1), we ultimately identified 6431 one-to-one orthologous genes for which we had sequences from all 14 investigated *Jaltomata* species and a unique tomato annotated coding sequence. All of these 6431 genes were used in the concatenation, majority rule, or quartet-based phylogeny reconstructions. From this dataset, we used 1517 genes in the BUCKy reconstruction (see Methods). Since we did not sample RNA from the reproductive tissues of *J*. *grandibaccata*, we excluded this species from analyses of locus-specific adaptive evolution; this resulted in a slightly larger number of orthologs, including those expressed solely in flowers. The resulting dataset had 6765 alignments of orthologous coding sequences, each containing sequences from the remaining 13 *Jaltomata* species (with *J*. *grandibaccata* excluded) and tomato. Among them, 4248 genes also have *C*. *annuum* orthologs, thus could also be used to test for positive selection on the ancestral branch leading to *Jaltomata*.

### Phylogenomic reconstruction of Jaltomata lineages supports several major clades

All four phylogenetic inference methods (concatenation, majority rule, quartet-based, and Bayesian concordance) generated a nearly identical species tree topology (Figure 2A, S3). In all trees, the first split in the species tree produces a well-supported clade that includes three north- and central-American species (*J*. *procumbens*, *J*. *repandidentata*, and *J*. *darcyana*) that all share floral traits (rotate corollas and light nectar) and make dark purple/purple fruit (CF=87). The remaining 11 species, that are found exclusively in South America and vary in floral traits and fruit colors, form a single moderately supported clade (CF=66). Based on sequence divergence at synonymous sites (Table S3), lineages within the non-purple-fruited clade have pairwise distance of 0.26% to 0.53%, and differ from the purple-fruited lineages by 0.95% to 1.29%. Within the nonpurple-fruited group, our reconstruction indicates that the red-fruited species *J*. *auriculata* is sister to the remaining species. The remaining species are split into a moderately supported clade (CF=67) of two species (*J*. *calliantha* and *J*. *quipuscoae*) that share floral traits and green fruits, and a relatively poorly supported clade (CF=19) consisting of the remaining eight species that vary extensively in floral traits but all produce orange fruit. Although quantitative support for different clades varied based on inference method (Figure S3), all trees produced the same topology, with the exception of inferred relationships within the orange-fruited group among *J*. *biflora*, *J*. *sinuosa*, *J*. *aijana* and *J*. *umbellata*. In that case, all concatenation, quartet-based, and concordance trees supported *J*. *sinuosa* as more closely related to *J*. *aijana* and *J*. *umbellata* (6% of gene trees support this grouping), while consensus trees placed *J*. *sinuosa* as sister to *J*. *biflora* (7% of gene trees support this grouping).

### Segregating variation is broadly shared among species in different subclades

To quantify how much variation shared among present subclades—presumably because of either shared ancestral variation or ongoing introgression—we mapped RNA-seq reads from each species to the tomato reference genome and called high-quality variants from ~8 million sites with more than 10X sequencing depth for all investigated species. We identified a large number of sites that are sorting the same allele among different subclades. Among them, 4303 variant sites are sorting in all four subclades (Figure 3A). We also quantified how many sites that are heterozygous in one lineage (accession) have the same two alleles sorting in other subclades. Within each lineage, the proportion of heterozygous sites range from 0.02%~0.16% (Table S4), which is comparable to the level of heterozygosity observed in self-compatible tomato species (Pease et al., 2016). For 13.57% to 64.35% of heterozygous sites in one species (Figure 3A, Table S4), both alleles could also be identified in other subclades, again indicative of a large amount of shared allelic variation.

**Fig 3.**
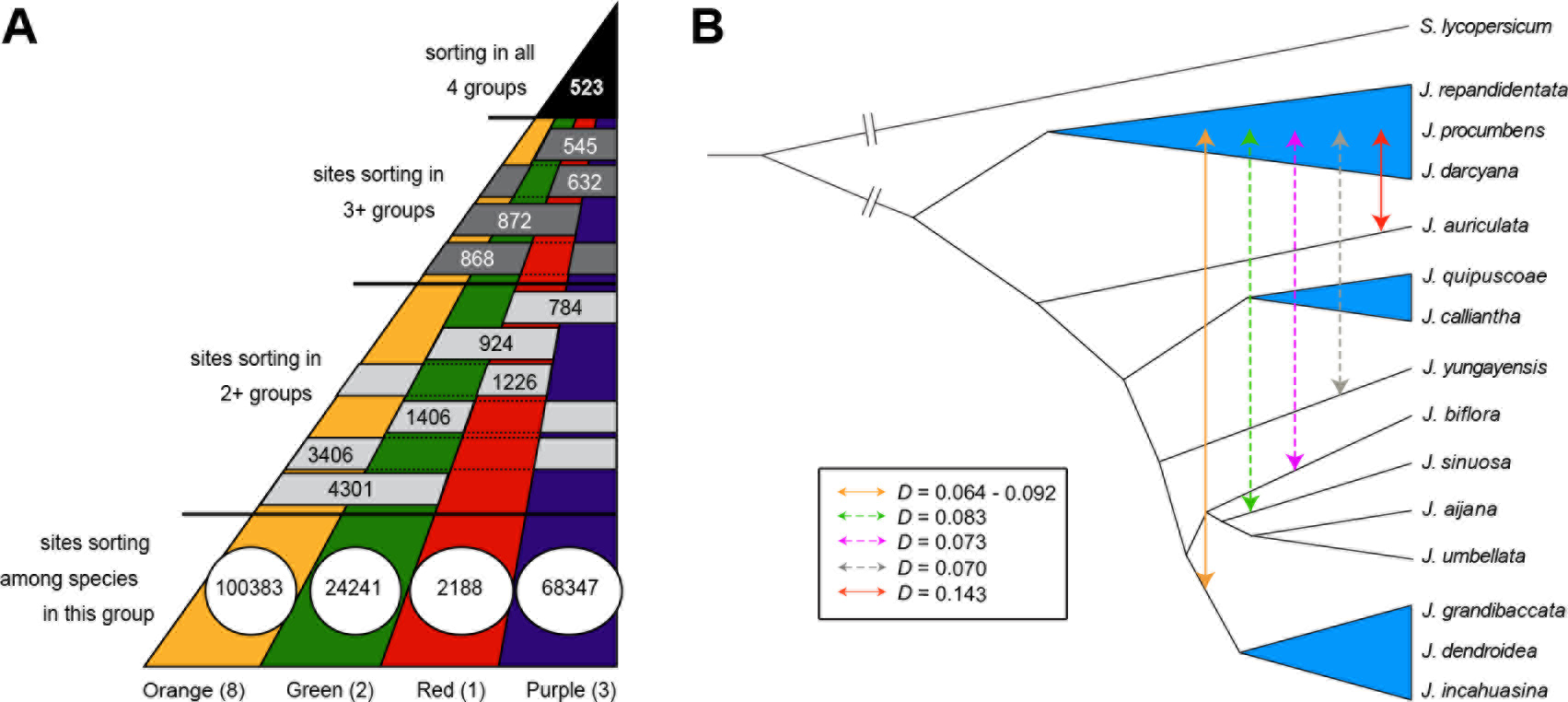
(A) Allele sorting, with the number of genetic variants within or shared between each *Jaltomata* subclade. (B) The introgression pattern among *Jaltomata* lineages. The solid lines indicate strong evidence of introgression between two lineages or sub-clades, while the dashed lines indicate putative introgression. The corresponding Patterson’s D statistic value is labeled for each putative introgression event.

### Phylogenomic discordance accompanies rapid diversification

As expected given the large number of genes (*n*=6431 and 6,223,350 sites in total) used for phylogenetic inference, our resulting species trees had very strong bootstrap support for almost all nodes (Figure S3A and S3C). Despite this, reconstructions also revealed evidence of extensive gene tree discordance consistent with rapid consecutive lineage-splitting events in this group (Figure 2B; S3B and S3D). For instance, the 6431 genes inferred 6431 different topologies, none of which matched the topology of the inferred species tree (Figure 2A). The concatenation tree has many extremely short internal branches, where gene trees show high levels of phylogenetic discordance. We detected a strong correlation between the internal branch length and levels of discordance (*P* = 0.0001, Figure S4), consistent with both ILS and introgression. Short branch lengths and extensive discordance were also detected for trees built individually for each of the 12 chromosomes (Figure S5; supplementary text).

Only three branches within *Jaltomata* are supported with relatively little discordance, i.e., with Bayesian CFs greater than 50 (Figure S3D): the branch leading to the purple-fruited clade, the branch uniting all non-purple-fruit *Jaltomata* lineages, and the branch leading to the two green-fruited lineages (Figure 2A). Along with the ancestral *Jaltomata* branch, these were the four branches on which most of our subsequent analyses were performed.

### Introgression after speciation among major clades of Jaltomata lineages

Given the apparent high level of phylogenetic discordance among our examined species, we tested for evidence of introgression on the background of presumed ILS. To do so, we calculated genome-wide *D*-statistics using the ABBA-BABA test. We only examined introgressions across well-supported subclades. In particular, we compared the distribution of trees in which one of two sister taxa (here, a species from either the red-, green-, or orange-fruited lineage) is closer to more distantly related species (in the purple-fruited clade) than the other. We found several such cases (Figure 3B). For example, there was a significant excess of sites that grouped the red-fruited lineage (*J*. *auriculata*) with a purple-fruited lineage, relative to the number of sites that grouped the green-fruited lineage with the purple-fruited lineage, indicative of detectable gene flow between the red-fruited and purple-fruited lineages since their split (Figure 3B and Table S5). We also inferred putative introgression, in at least two separate events, involving six species in the orange-fruited clade with the purple-fruited clade (Figure 3B and Table S5). First, we inferred a shared introgression event between the purple-fruited group and three of the orange-fruited species (*J*. *grandibaccata*, *J*. *dendroidea*, and *J*. *incahuasina*); this excess includes shared specific sites that support the same alternative tree topology for each of these three ingroup species, suggesting that it likely involved the common ancestor of all three contemporary orange-fruited species (Table S6). Second, we detected evidence for gene flow between the remaining orange-fruited species (*J*. *yungayensis*, *J*. *biflora* and *J*. *sinuosa*) and the purple-fruited lineage, in the form of significant genome-wide *D*-statistics (Table S5). Because we did not observe an excess of shared specific sites supporting the same alternative tree topology among these three orange-fruited species (Table S6), these patterns are suggestive of three putative independent introgression events. However, given very low resolution of patterns of relatedness among orange-fruited species, the specific timing of these events is hard to resolve.

### Ancestral state reconstruction suggests different histories for fruit color and floral trait evolution

Based on the inferred species tree (Figure 2A), we reconstructed the ancestral states of fruit and floral traits (Figure 4; S6). The four subclades of *Jaltomata* species were inferred to have evolved different fruit colors at their corresponding common ancestors (Figure 4A). Our reconstruction suggests that the derived nectar traits (orange/red nectar color, and increased nectar volume) probably evolved at the common ancestor of the green/orange-fruited clade (Figure S6A and S6B), with two subsequent reversions to ancestral conditions within this clade. The evolution of the two derived corolla shapes in *Jaltomata* (campanulate and tubular) appears to be more complex (Figure 4B). At the majority of internodes within the non-purple-fruited lineages, all three corolla shapes (i.e. rotate (ancestral), campanulate and tubular) show ≥10% probability of being the ancestral state, making specific inferences about corolla shape evolution within this clade uncertain. Consistent with these patterns, we found that concordance factors were very low at almost all internodes within the radiating subgroup that displays the derived floral traits (i.e., the non-purple-fruited lineages) (Figure 2A; S3D), whereas they were considerably higher on branches associated with fruit color evolution (including the branch uniting the two green-fruited species analyzed).

**Fig 4.**
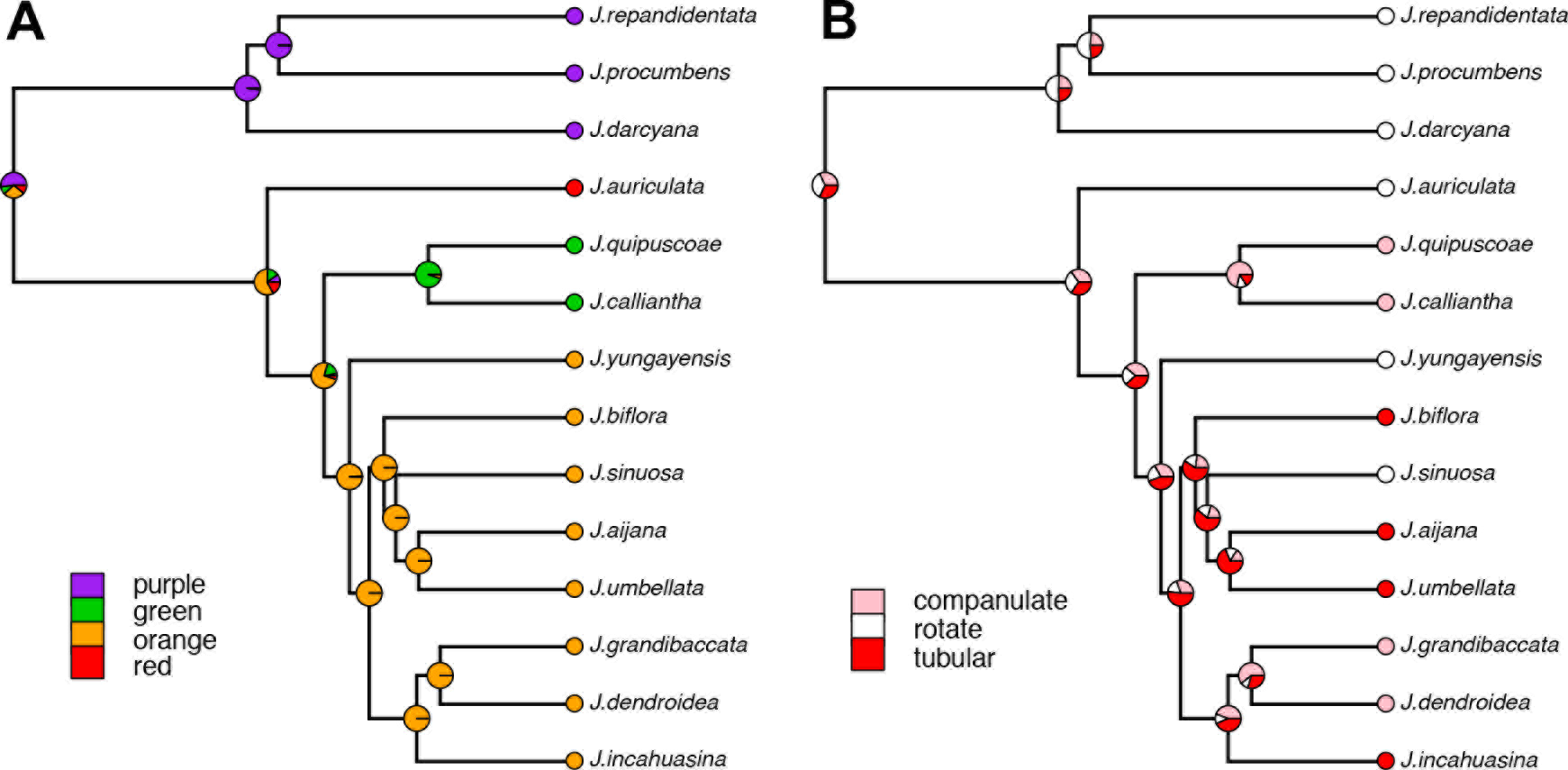
Ancestral character-state reconstruction of **(A)** fruit color, **(B)** corolla shape in investigated *Jaltomata* using maximum likelihood.

These analyses suggest alternative evolutionary and genetic histories for our traits of interest. In particular, strong associations between fruit color transitions and specific branches/clades within *Jaltomata* suggests that the underlying genetic changes are more likely due to conventional lineage-specific *de novo* evolution along the relevant branches. In contrast, the distribution of floral trait variation produces an ambiguous reconstruction of trait transitions, especially for floral shape, such that the distribution of ancestral versus derived floral shape variation is unassociated with phylogenetic relationships in the non-purple-fruited *Jaltomata* clade (Figure 4B). While strictly *de novo* evolution occurring multiple times is not excluded as an explanation of floral evolutionary transitions, one alternative is that these trait transitions drew upon shared variation segregating in the ancestor of these lineages. Accordingly, in the next sections we evaluate both lineage-specific *de novo* evolution and selection from standing ancestral variation when searching for genetic variants that might have contributed to floral trait evolution. The lineage-specific *de novo* evolution analysis alone is used to identify potential candidates for fruit color evolution, since the approach we use to identify standing ancestral variation is only informative in cases where trait variation is not confounded with phylogenetic relationship.

### Loci with patterns of positive selection associated with lineage-specific trait evolution

We performed tests of molecular evolution for all orthologous clusters that contained a sequence from every *Jaltomata* accession and an ortholog from the tomato outgroup. Depending upon the specific branch being tested, we detected evidence for positive selection in ~1-2% of loci in our dataset, based on whether the locus had significantly elevated *d*_*N*_/*d*_*S*_ ratios (*d*_*N*_/*d*_*S*_ >1; *p* < 0.01). This included 1.88% of genes (67 out of 3556 testable genes; Table S7) on the *Jaltomata* ancestral branch, 1.58% in the purple-fruited group (48 out of 3033 testable genes; Table S8), 2.61% in the red-fruited group (70 out of 2686 testable genes; Table S9), 1.96% in the green-fruited group (30 out of 1531 testable genes; Table S10), and 0.74% in the non-purple-fruited *Jaltomata* lineages (15 out of 2039 testable genes; Table S11). Many of the genes showing elevated *d*_*N*_/*d*_*s*_ appear to have general molecular functions (e.g. transcription, protein synthesis, or signaling), including numerous genes involved in various stress responses, such as heavy metal tolerance, sugar starvation response, protection from ultraviolet (UV) radiation and extreme temperature, and herbivore and pathogen resistance (Table S7-11). Our positively selected loci contain genes functionally associated with photosynthesis, fatty/lipid biosynthesis and transportation, and sugar signal transduction (observed in the GO enrichment analysis; Table S12-16), as well as loci with unknown functions.

After controlling for multiple tests using an FDR <0.1 on each branch tested, only three genes on the *Jaltomata* ancestral branch, one gene on the purple-fruited ancestral branch, and four genes on the red ancestral branch, remained significant for *d*_*N*_/*d*_*S*_>1. Interestingly, for 5 of these 8 loci, the inference of positive selection appears to be due to the presence of a multi-nucleotide mutation (MNM) specifically on the target branch, a mutational pattern known to produce spurious inferences of positive selection in PAML’s branch-site test (Venkat et al., 2017). With one exception, all of these potential false positives were found on the ancestral *Jaltomata* or purple-fruited clade branches, the longest Jaltomata-specific branches in our analyses; as MNMs are more likely to appear on long branches (they are relatively rare mutations) these are expected to be enriched for these spurious inferences in the branch-site test (Venkat et al., 2017). Our three remaining loci, with positive selection on the red-fruited branch, include a gene (*BANYULS*; ortholog to *Solyc03g031470*) with functional roles in pigmentation (see Discussion).

We also detected several instances where slightly less stringent criteria (*d*_*N*_/*d*_*S*_ >1; *p* <0.05) revealed lineage-specific adaptive evolution of our *a priori* candidate genes occurring on a branch that is also inferred to be associated with the evolution of derived traits (Table S17). Most notably, we found evidence of positive selection on candidate loci that are likely to be involved in fruit color, including a gene encoding *ζ*-carotene isomerase (*Z-ISO*; ortholog to *Solyc12g098710*) on the red-fruited lineage, and the two other genes significant on the ancestral branch of the green-fruited lineages encoding carotenoid cleavage enzyme 1A (*CCD1A*; ortholog to *Solyc01g087250)* and zeaxanthin epoxidase (ZEP; ortholog to *Solyc02g090890*) (Figure 5, see Discussion). We detected signatures of positive selection on fewer of the genes involved in floral development, mostly notably in the MADS-box gene *APETALA3* (*AP3*/*DEF*, ortholog to *Solyc04g081000*) on the ancestral branch to the purple-fruited lineage. Overall, we note that many of our loci (including *a priori* candidates) did not meet the requirements to be tested for positive selection (Table S7); in particular, gene trees for many loci lacked the required support for a specific internal branch, either because of incongruence or limited phylogenetic signal, especially within the rapidly diverging orange-fruited clade (Table S17).

**Fig 5.**
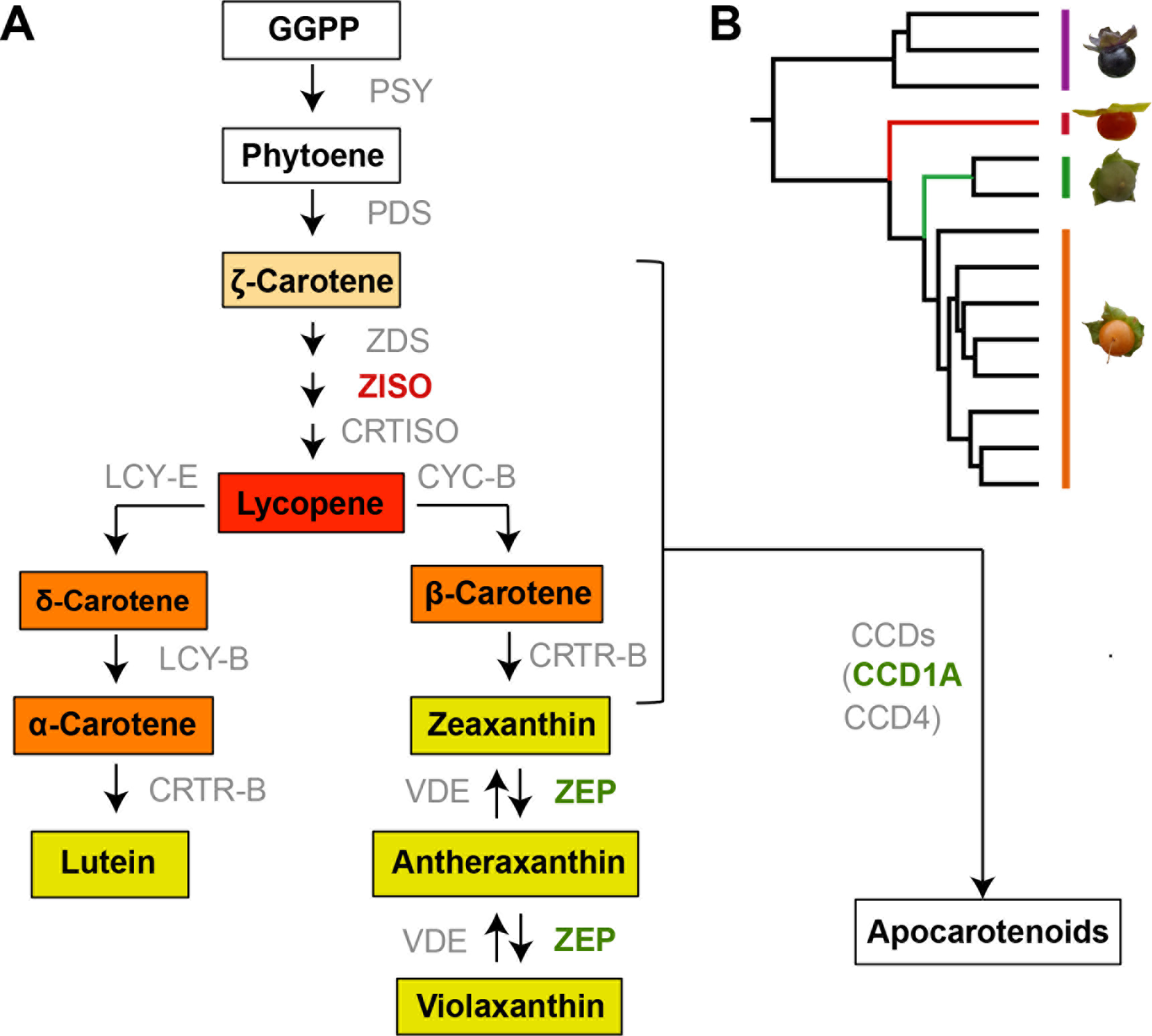
Genes under adaptive evolution in the carotenoid biosynthesis pathway. **(A)** Simplified carotenoid biosynthesis pathway modified from (Yuan et al. 2015). Genes under adaptive evolution are indicated by their names highlighted in colors corresponding to particular branches (see panel B). *PSY*, phytoene synthase; *PDS*, phytoene desaturase; *ZDS*, *ζ*-carotene desaturase; *ZISO*, *ζ*-carotene isomerase; *CRTISO*, carotenoid isomerase; *LCY-E*, lycopene *ε*-cyclase; *LYC-B*, lycopene β-cyclase; *CRTR-B*, *β*-ring hydroxylase; *CYC-B*, chromoplast specific lycopene *β*-cyclase; *ZEP*, zeaxanthin epoxidase; *VDE*, violaxanthin de-epoxidase; *CCD*, carotenoid cleavage dioxygenase. Metabolites are boxed and colored according to their compound colors, whereas white boxes indicate no color. **(B)** Positive selection signatures of genes on different branches are indicated by different colors: red-fruited lineages (Red), and green-fruited lineages (Green).

### Loci potentially associated with trait evolution from standing ancestral variation

To investigate whether ancestral variants are potentially associated with floral trait diversification, we performed a “PhyloGWAS” analysis (Pease et al., 2016). Such variants will have differentially fixed among descendant lineages, leading to genes that cluster species together based on floral traits regardless of their overall phylogenetic relationships. We found 31 genes with nonsynonymous variants perfectly associated with the derived floral traits (Table S18), which was significantly more than the number of loci expected by chance to have segregation patterns that exactly match the tip states (*p* < 9.3 × 10^-5^). Most of these genes are characterized by only one or few nucleotide differences, which is an expected pattern for variants recently selected from standing ancestral variation (Pease et al., 2016). These results suggest that one or few molecular variants present in ancestral populations could contribute to the multiple apparent transitions to derived floral shapes in *Jaltomata*. Among the loci identified by our approach, some genes are potentially functionally related to petal development, including *ARGONAUTE1* (*AGO1*) and xyloglucan endotransglucosylase/hydrolase 2 (*DcXTH2*) (see Discussion).

## DISCUSSION

Within rapidly radiating groups, the patterns of genetic relatedness among lineages provide essential data for determining the pace and location of important trait transitions, and their underlying genes; both are critical for understanding the drivers of rapid diversification and speciation. Our phylogenomic analyses of the 14 investigated *Jaltomata* species revealed genome-wide gene tree discordance, and a highly complex history of genetic relatedness among contemporary lineages, consistent with other studies of recently radiating groups (Brawand et al., 2014; Garrigan et al., 2012; Novikova et al., 2016; Pease et al., 2016). We identified substantial ILS and shared ancestral polymorphism, as well as evidence of putative introgressions among the subclades of *Jaltomata* species, as the sources of this observed complex genome-wide history. This complexity was also reflected in inferences about the evolution of major trait transitions within the group. We found differences in the patterns of fruit versus floral character evolution and in our inferred confidence in the reconstruction of these patterns, including their likely risk of hemiplasy. Given this, we used several strategies to identify loci that might contribute to the evolution of these traits, including examining lineage-specific *de novo* adaptive evolution along well-supported branches and identifying variants that might have been selected from standing ancestral variation. Overall, by combining evidence from molecular evolution with data on trait variation across a clade—and a more direct accounting for the risk of hemiplasy—we generated more conservative, but credible, inferences of candidate genes responsible for the evolution of ecologically important phenotypic traits. Specifically, we identified several functionally relevant candidate genes for our target trait transitions, and different potential sources of adaptive evolution fueling changes in target floral versus fruit traits.

### Extensive ILS and several introgression events produce a complex genome-wide history during rapid diversification

Our reconstruction of phylogenetic relationships based on a transcriptome-wide dataset agrees with previous studies that identified three major *Jaltomata* sub-clades primarily distinguished from each other by their fruit colors (Miller et al., 2011; Särkinen et al., 2013), including the previous inference that the clade of purple-fruited species is sister to the rest of the genus (Särkinen et al., 2013). However, the relationships among species within each subclade show high levels of discordance, and the distribution of genetic variation strongly indicates a rapid and recent evolutionary origin; the internal branch lengths within each subclade are short, especially in the non-purple-fruited lineages (Figure 2A). Accordingly, the history of these species—including the orange/green-fruited lineages that show the most floral trait diversity among *Jaltomata* species—is expected to be strongly affected by ILS and introgression.

Indeed our genome-wide reconstruction indicated that, although concatenation or quartet-based approaches generated a species tree with high bootstrap support values (commonly observed when inferring trees from large amounts of data (Kubatko & Degnan, 2007; Salichos & Rokas, 2013), gene tree discordance was rampant, and individual gene trees showed highly variable support for the specific placement of individual species (Figure 2B) especially at short branches (Figure 2A). Our finding of extensive genome-wide ILS in the genus *Jaltomata* agrees with other recent studies on contemporary (Novikova et al., 2016; Pease et al., 2016) and relatively ancient adaptive radiations in plant species (Wickett et al., 2014; Yang et al., 2015), and is emerging as a universal signal of rapid radiation in comparative genome-wide datasets.

Resolution of phylogenetic relationships among the species within each *Jaltomata* subclade was insufficiently clear to investigate introgression within subgroups. However, across major sub-clades we identified at least two clear introgression events that involved either orange and red-fruited lineages with the purple-fruited lineages, similar to the detection of introgression events in other recent genome-wide studies on closely related plant species (Eaton & Ree, 2013; Novikova et al., 2016; Owens et al., 2016; Pease et al., 2016). Interestingly, in one case an excess of sites supported a shared introgression pattern between three orange-fruited species (i.e. *J*. *grandibaccata*, *J*. *dendroidea*, and *J*. *incahuasina*) and the purple-fruited clade, consistent with a scenario in which introgression involved the recent common ancestor of these three orange-fruited species. Moreover, in this case, the inference of a single shared introgression event itself provided more confidence in this specific ancestral branch within the orange-fruited clade. Overall, as with ILS, post-speciation introgression is another inference frequently emerging from contemporary phylogenomic studies of radiations.

### Inferring the history of trait evolution and the contributing loci in the presence of rampant discordance

The complex history of genomic divergence in this clade has clear consequences for inferences of trait and gene evolution. When ILS or introgression can plausibly explain the discordant distribution of traits, it might be impossible to infer trait evolution with any certainty in the absence of additional independent information about target traits, such as their genetic basis (Hahn & Nakhleh, 2016). In contrast, the evolutionary transitions of some traits might be confidently inferred as long as the relevant branches and resulting relationships are associated with higher levels of concordance (Hahn & Nakhleh, 2016). Species in *Jaltomata* exhibit extensive trait diversity, most notably in fruit color, corolla shape, and nectar volume and color (Figure 2A) (Miller et al., 2011), and one of the main goals in this study was to better understand the evolutionary history of these trait transitions, including the genetic basis of the traits associated with rapid phenotypic diversification. A previous phylogenetic study based on a single locus suggested that floral traits (including corolla shape and nectar color) might have evolved multiple times independently in *Jaltomata* species (Miller et al., 2011). However, the presence of rampant discordance, and abundant evidence of shared ancestral variation, makes inferring the history of trait transitions and their genetic basis especially challenging in this group. Indeed, our analyses indicated that different classes of trait transition—most notably fruit color versus floral shape variation—were differently susceptible to hemiplasy. For floral shape evolution in *Jaltomata*, a lack of resolution and high gene tree discordance at key nodes within the phylogeny, including within the clade displaying the greatest phenotypic diversity (Figure 2A and Figure 4B), mean that hemiplasy is a plausible explanation of the discordant distribution of similar floral traits—rather than multiple independent evolutionary events (i.e. homoplasy). In contrast, we showed that the history of fruit color evolution could be confidently inferred, as these trait transitions occurred on branches with higher levels of concordance and therefore lower risks of hemiplasy (Figure 2A and Figure 4A).

Accounting for the potential influence of hemiplasy is also critical when generating hypotheses about the loci that could have contributed to trait transitions. In general, incorrect reconstructions of trait history will suggest incorrect candidates involved in the evolution of those traits. Moreover, tests of molecular evolution can be specifically misled if trait transitions occur on discordant gene trees (Mendes et al., 2016). Accordingly, to identify loci that might be responsible for any particular trait transitions, different approaches will be appropriate depending upon the confidence with which hemiplasy can be excluded or not. For traits evolving on branches where discordance is low, confidence is high, and hemiplasy is unlikely, it is reasonable to expect that lineage-specific *de novo* substitutions are a substantial contributor to relevant trait evolution. Given a high risk of hemiplasy, genetic variation underpinning trait evolution could potentially come from additional sources, including recruitment of ancestral polymorphisms and/or introgression. These differences are exemplified in our study by the alternative histories, and different genetic hypotheses, generated for fruit color versus floral shape traits.

**Floral shape evolution** – Based on the inferred species tree (Figure 2A), the 14 investigated *Jaltomata* species are not related according to their floral shape traits (i.e. floral shape is distributed paraphyletically). Moreover, the branch lengths leading to lineages with derived character states are uniformly short with high levels of gene tree discordance (Figure 2A), so that the probability of hemiplasy is expected (Hahn & Nakhleh, 2016) to be very high. Indeed, when we reconstruct the evolution of corolla shape (Figure 4B), the three alternative corolla morphs were inferred to be almost equally likely at the common ancestor of non-purple-fruited lineages. It is possible that this lack of resolution is due to introgression. For example, the ancestral floral character states (i.e. rotate corolla shapes and clear nectar) found in *J*. *yungayensis* and *J*. *sinuosa* within the orange-fruited clade could be due to alleles introgressed from purple-fruited species, as we identified putative introgression events between those lineages (Figure 3B). However, the lack of a reference genome for *Jaltomata* precluded us from more directly investigating evidence (for example, locus-specific patterns of introgression) that introgression might contribute to the distribution of ancestral floral traits within this sub-clade. Instead, because of the high risk of hemiplasy and low resolution of ancestral states, we used several approaches to identify the genetic variants associated with the two other potential sources of trait variation.

First, if the paraphyletic distribution of derived traits is due to hemiplasy among species, the relevant nucleotide differences should be at the same sites in all lineages that share derived traits (Hahn & Nakhleh, 2016). Using this rationale, Pease et al. (2016) identified tens to hundreds of genetic variants among wild tomato lineages that were exclusively associated with each of three ecological factors, variants that are candidate targets of parallel ecological selection on standing ancestral variation in that group. Here, we used an analogous approach to look for variants associated with phenotypic (floral) trait variation in *Jaltomata*, and identified 31 candidate genes with nonsynonymous variants that are completely correlated with the distribution of floral trait variation (derived vs. ancestral; Figure S2). Among them, the gene *AGO1* (ortholog to *Solyc02g069260*) is known to be necessary in *Arabidopsis* floral stem cell termination and might act through *CUC1* and *CUC2* (*a priori* candidate genes), which redundantly specify boundaries of floral meristem (Ji et al., 2011; Kidner & Martienssen, 2005). Another gene *DcEXPA2* (ortholog to *Solyc02g091920*) is known to be markedly up-regulated in the petals of carnation (*Diathathus caryophyllus*), and is potentially associated with the petal growth and development (Harada et al., 2010). We also note that, although these shared hemiplasious variants could be due to post-speciation introgression rather than sorting from ancestral variation, given the often allopatric geographical distribution of our lineages (Figure 1) and evidence of substantial allelic sharing among contemporary subclades (Figure 3A), we infer that these variants were more likely sorted from ancestral variation.

Second, because our reconstruction of floral trait evolution could not exclude the role of lineage-specific *de novo* mutation, we also examined our transcriptomes for genes showing lineage-specific evolution associated with the derived floral traits. However, we found that only a small number of genes were testable due to the extensive gene tree discordance specifically at the branches leading to subclades with the derived floral characters. Moreover, none of our *a priori* candidate genes involved in floral development showed suggestive patterns of molecular evolution on internal branches of *Jaltomata* (Table S7), nor did we find any other functionally suggestive (floral development-related) genes adaptively evolving on branches leading to specific subclades within *Jaltomata*, with the exception of *APETALA3* (*AP3*)—a gene associated with the formation of petals and stamens in flowering plants, including *Arabidopsis*(Wuest et al., 2012; Specht & Howarth, 2015)—although this was detected on the branch leading to the purple-fruited clade, within which all species retain the ancestral rotate corolla form.

***Fruit color evolution*** – In contrast to floral traits, ancestral state reconstruction in the 14 investigated *Jaltomata* species suggested that fruit color transitions follow phylogenetic relationships. The ancestral states of fruit colors at most relevant internodes can be inferred with high confidence (Figure 4A), and the internal branches leading to fruit trait transitions are well-supported, indicating a low probability of hemiplasy (Figure 2A). Accordingly, we identified a set of loci showing adaptive evolution specifically on these branches, provided that each tested gene tree 1) supported the internal branch being tested, and 2) showed at least one clade-specific nonsynonymous substitution on that branch (Mendes et al., 2016; Pease et al., 2016), to avoid potential inference errors associated with examining genes that have topologies discordant with the species tree (Mendes et al., 2016). Our analyses revealed multiple adaptively evolving candidate loci with clear functional relevance to these trait transitions, including several *a priori* candidate genes involved in the carotenoid pathway (Yuan et al., 2015) (Figure 7). First, *Z-ISO* (ortholog to *Solyc12g098710*), a key enzyme in the production of red-colored lycopene in the carotenoid biosynthetic pathway (Chen et al., 2010), was positively selected on the branch leading to our one species with bright red fruit, *J*. *auriculata*. Interestingly, this gene was also previously found to show adaptive evolution specifically on the branch leading to the red-fruited (Esculentum) group in wild tomatoes (Pease et al., 2016). Second, on the branch leading to our two green-fruited species, we found significantly elevated *d*_*N*_/*d*_*S*_ ratios for both *CCD1A* (ortholog to *Solyc01g087250*), a gene whose product participates in the conversion of carotenoid pigments to isoprenoid volatiles (Ilg et al., 2014), and for *ZEP* (ortholog to *Solyc02g090890*), which converts zeaxanthin to violaxanthin (Marin et al., 1996). *CCD1* has previously been identified in tomato fruits as responsible for generating flavor volatiles (Auldridge et al., 2006; Simkin et al., 2004). This functional observation from a closely related group is intriguing because fruits of our two green-fruited species (*J*. *quipuscoae* and *J*. *calliantha*) appear to produce the strongest scent within the 14 *Jaltomata* species analyzed here (J. Kostyun, unpubl. data). This apparent increase in fragrance—presumably due to changed volatile organic compounds—might play a role in attracting vertebrate frugivores for seed dispersal.

In addition to carotenoids, among *a priori* candidate genes involved in the biosynthesis pathway of water-soluble vacuolar anthocyanin pigments, we detected *BANYULS* (ortholog to *Solyc03g031470*) selected on the red-fruited branch. (Note that PAML also indicates adaptive molecular evolution of this locus on the purple-fruited branch, but this appears to be due to the presence of a MNM; see results). In addition to *a priori* candidates, genome-wide unbiased analyses also identified that multiple genes belonging to *R2R3MYB*, *BHLH* and *WD40*-repeats classes of loci were under positive selection in purple-fruited lineages and the red-fruited lineage. The *MYB-BHLH-WD40* TF complexes are known to regulate cellular differentiation pathways, including of the epidermis, as well as transcription of anthocyanin structural genes (Gonzalez et al., 2008; Jaakola, 2013; Ramsay & Glover, 2005).

### Implications for the inference of phenotypic trait evolution and causal genetic variation in rapid radiating lineages

As highlighted here and in numerous recent phylogenomic studies (Eaton & Ree, 2013; Novikova et al., 2016; Owens et al., 2016; Pease et al., 2016), both ILS and introgression contribute to the history of diversification within radiating clades, so that evolution in these groups is more complex than can be represented by a simple bifurcating species tree. This complexity has important implications for empirical inferences about historical relationships and trait evolution, because assuming resolved relationships without taking into account incongruence can fundamentally mislead inferences in both these cases (Hahn & Nakhleh, 2016). It also must be accounted for when considering the genetic changes that might have fueled diversification; when a trait has several possible alternative evolutionary histories, it is necessary to investigate the range of alternative sources of genetic variation—including *de novo* lineage-specific evolution and selection from ancestral variation—that could fuel this trait evolution. Here, we provided a genome-wide analysis of the recently diversified plant genus *Jaltomata* in which we consider the relative risk of hemiplasy while identifying candidates for the specific loci underlying trait evolution. Our analysis highlights a growing appreciation that rapid radiations can and likely do draw on multiple sources of genetic variation (Hedrick, 2013; Pease et al., 2016; Richards & Martin, 2017). Indeed, while independently originating variants could explain the recurrent evolution of phenotypic similarity—a frequent observation in adaptive radiations—it is clear that shared ancestral genetic variation, or alleles introgressed from other lineages, have also made substantial contributions (Elmer & Meyer, 2011; Stern, 2013). Going forward, it will be necessary to distinguish between these alternative scenarios to understand how different evolutionary paths contribute to phenotypic convergence and differentiation (Martin & Orgogozo, 2013; Stern, 2013) and to identify the specific variants responsible.

## ACKNOWLEDGEMENTS

The authors thank James Pease, Rafael Guerrero and Fabio Mendes for advice on performing comparative phylogenomic and molecular evolution analyses. This research was funded by National Science Foundation grant DEB-1135707, to LCM and MWH.

## AUTHOR CONTRIBUTIONS

L.C.M, M.W., M.W.H., and J.L.K. designed the experiments; J.L.K generated the experimental materials; M.W. conducted the bioinformatics analyses; and M.W. and L.C.M. wrote the paper with contributions from M.W.H. and J.L.K.

## DATA AVAILABILITY

Raw reads (FASTQ files) for generating the 14 species transcriptomes are deposited in the NCBI SRA (BioProject: PRJNA380644). Multiple sequence alignments used for phylogenetic tree reconstruction and molecular evolution analyses are available at Dryad (doi: XXX). All commands and scripts used for analyses in this study can be found in our project directory on GitHub (https://github.com/wum5/JaltPhylo).

